# Repeated unilateral handgrip contractions alter functional connectivity and improve contralateral limb response times: A neuroimaging study

**DOI:** 10.1101/2021.10.23.464569

**Authors:** Justin W. Andrushko, Jacob M. Levenstein, Catharina Zich, Evan C. Edmond, Jon Campbell, William T. Clarke, Uzay Emir, Jonathan P. Farthing, Charlotte J. Stagg

## Abstract

In humans, motor learning is underpinned by changes in sensorimotor network functional connectivity (FC). Unilateral contractions increase FC in the ipsilateral primary motor cortex (M1) and supplementary motor area (SMA); areas involved in motor planning and execution of the contralateral hand. Therefore, unilateral contractions are a promising approach to augment motor performance in the contralateral hand. In a within-participant, randomized, cross-over design, 15 right-handed adults had two magnetic resonance imaging (MRI) sessions, where functional-MRI and MR-Spectroscopic Imaging were acquired before and after repeated right-hand contractions at either 5% or 50% maximum voluntary contraction (MVC). Before and after scanning, response times (RTs) were determined in both hands. Nine minutes of 50% MVC contractions resulted in decreased handgrip force in the contracting hand, and decreased RTs in the contralateral hand. This improved motor performance in the contralateral hand was supported by significant neural changes: increased FC between SMA-SMA and increased FC between right M1 and right Orbitofrontal Cortex. At a neurochemical level, the degree of GABA decline in left M1, left and right SMA correlated with subsequent behavioural improvements in the left-hand. These results support the use of repeated handgrip contractions as a potential modality for improving motor performance in the contralateral hand.

**Significance Statement:** Enhanced functional connectivity and decreased inhibition in sensorimotor areas of the brain underpin enhancements in motor performance and learning. In this study we investigated the impact of repeated right handgrip contractions at 50% MVC to improve behaviour, enhance functional connectivity and alter sensorimotor inhibition. We found that after nine minutes of repeated 50% MVC handgrip contractions with the right-hand, left-hand response times were significantly improved. This behavioural improvement was accompanied by altered interhemispheric functional connectivity and neurochemical changes across sensorimotor areas. Repeated unilateral handgrip contractions may be an effective method for enhancing contralateral limb motor performance in rehabilitation settings.

## Introduction

The acquisition of motor skill is essential in our daily lives. Able-bodied individuals may take for granted the abilities to grasp and manipulate objects, or even to safely complete essential activities of daily living. However, many people lose their ability to perform these tasks through a variety of injuries. Restoring function in these individuals is of clear clinical importance, but how we can optimally improve behaviour is an open scientific question: both in terms of restoring motor function in orthopedic or neurologically impaired individuals such as stroke survivors, and in healthy populations, such as athletes, looking to maximize motor performance.

At a systems level, studies suggest that motor performance can be enhanced through interventions that either increase corticomotor excitability (Sanes & Donoghue, 2000; Cirillo *et al*., 2010, 2011; Lissek *et al*., 2013) and/or decrease corticomotor inhibition. Gamma-aminobutyric acid (GABA)-ergic inhibition is shown to decrease during motor learning (Floyer-Lea *et al*., 2006; Kolasinski *et al*., 2019), and the magnitude of this GABA decrease correlates with the extent of subsequent motor performance improvement (Stagg *et al*., 2011*a*). Altering motor cortex (M1) excitability and inhibition through non-invasive brain stimulation techniques increases motor skill acquisition, both in neurologically intact adults (Reis *et al*., 2009; Stagg *et al*., 2011*b*) and in functional recovery after stroke (Kim *et al*., 2010; Allman *et al*., 2016).

However, non-invasive brain stimulation is not widely available, and determining other methods to alter corticomotor function is an important research objective. One approach that is widely accessible and has shown promise for altering the cortical excitation/inhibition balance is performing a unilateral exercise protocol that induces motor fatigue. At a neurochemical level, unilateral fatiguing exercise decreases M1 GABA_A_ activity, as quantified by the Transcranial Magnetic Stimulation protocol Short-interval Intracortical Inhibition (SICI) in the contralateral M1 (cM1) (Maruyama *et al*., 2006) and ipsilateral M1 (iM1) (Takahashi *et al*., 2009). Unilateral fatiguing exercise also increases iM1 cortical excitability (Aboodarda *et al*., 2016, 2019), and functional connectivity in the sensorimotor network, particularly in the iM1, ipsilateral supplementary motor area (SMA), and contralateral SMA (Jiang *et al*., 2012). Unilateral fatiguing exercise has been shown to enhance muscle activity in the contralateral unfatigued homologous muscle (Carr *et al*., 2021), although this finding is not consistent across the literature (Kavanagh *et al*., 2016; Branscheidt *et al*., 2019). Here, we wished to understand whether these ipsilateral cortical alterations serve as a neural basis for contralateral behavioural improvements.

Considering motor network connectivity is at least in part controlled by M1 inhibition (Stagg *et al*., 2014; Bachtiar *et al*., 2015), we acquired two independent measures [resting-state functional magnetic resonance imaging (rs-fMRI), and resting-state magnetic resonance spectroscopic imaging (rs-MRSI)]) to test the hypothesis that repeated unilateral handgrip contractions, resulting in improved performance in the opposite hand (i.e., faster response times) are related to increased interhemispheric homologous connectivity of M1, and SMA via increased glutamate and/or decreased GABA in these regions (figure 1A).

**Figure 1.**
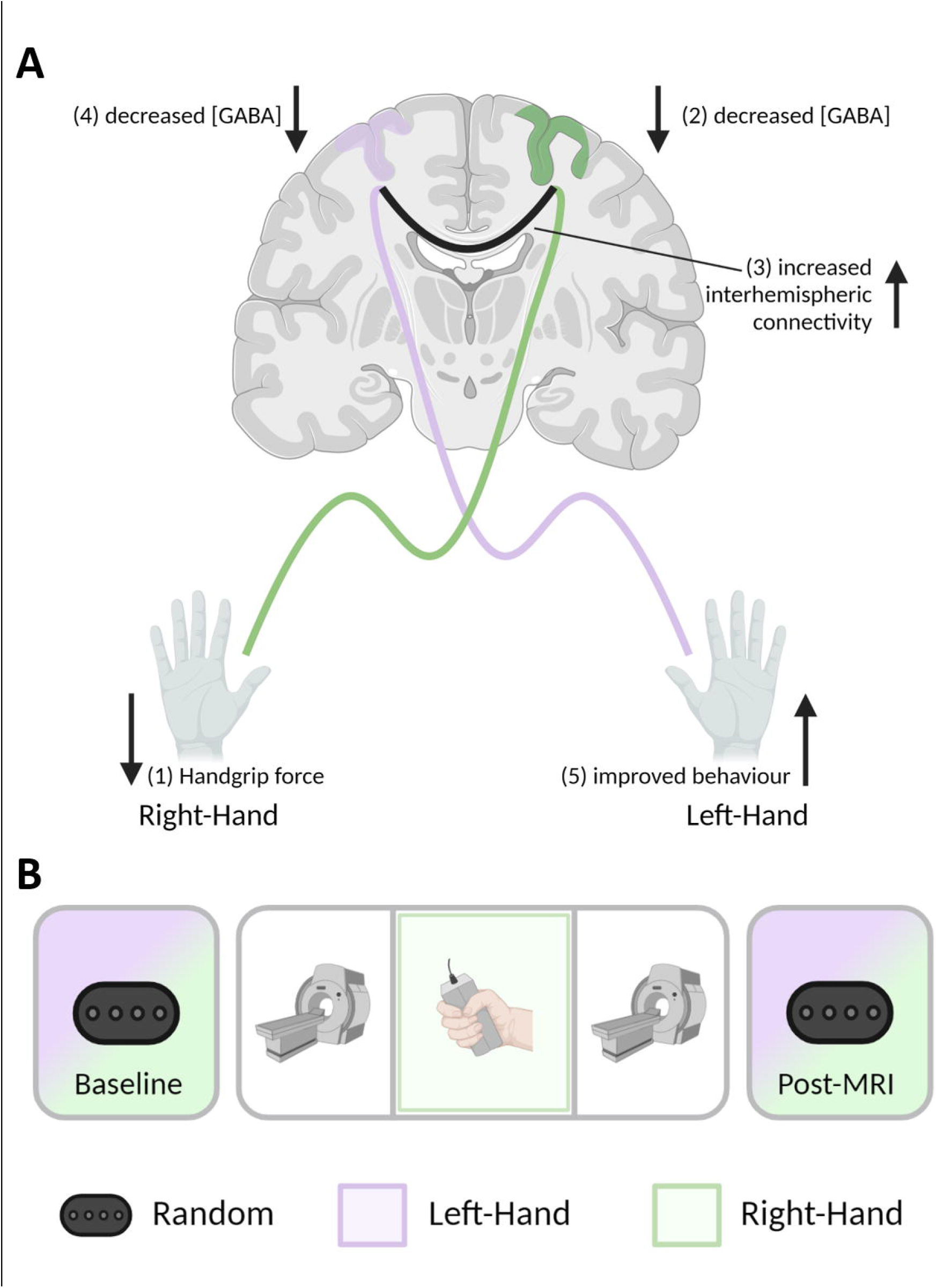
**A)** Theoretical physiological model for the present study, **B)** Schematic outlining the study design. Both aspects of this figure were created with BioRender.com.

## Methods

This study conforms to the Declaration of Helsinki and was approved by the Oxford Central University Research Ethics Committee (MSD-IDREC-C1-2014-100 and MSD-IDREC-C1-2014-090). Using an effect of η^2^ = 0.250 based on the relevant interaction from a previous brain stimulation study (transcranial direct current stimulation: anodal, cathodal, bilateral sham) × neurochemical (GABA, glutamate) × hemisphere (Left M1, right M1) × time (during stimulation, post 1, post 2, post 3) (Bachtiar *et al*., 2018). This higher-order interaction was chosen given the similarity in study design of a unilateral protocol designed to induce interhemispheric interactions at a neurochemical level. We computed an estimated total sample size of 12 (G*Power 3.1.9.2; 1-β = 0.95, α = 0.05). To allow for dropout and data loss, we therefore recruited fifteen right-handed participants [Edinburg Handedness Questionnaire – short form (EHQ) (Oldfield, 1971; Veale, 2014)] aged between 18-35 years (28.92 ± 3.30 years; 7 Female; EHQ: 85.8 ± 18.8). Exclusion criteria were: playing a musical instrument to Associated Board of the Royal Schools of Music Grade 5 or higher, any contraindication to MRI, any history of neurological or psychiatric disorders.

### Experimental Outline

Participants had two experimental sessions where they performed a right handgrip force-matching task at either 5% or 50% of their maximum voluntary contraction (MVC) in a randomized order (n = 7 started with 5% MVC; n = 8 started with 50% MVC). During each experimental session, response times (RTs) of each hand were assessed before and after the MRI. Resting-state functional MRI (fMRI) and magnetic resonance spectroscopic imaging (MRSI) scans were acquired before and after a nine-minute right handgrip force-matching task at either 5% or 50% MVC. This study had a within-participant repeated measures design, with the order of conditions stratified-randomized for sex across the group (figure 1B).

### Handgrip force-matching task

The handgrip force-matching task was performed using an MRI-compatible hand dynamometer (Biopac Systems Inc. Aero Camino Goleta, CA) and implemented using in-house code (Psychtoolbox-3 (Kleiner *et al*., 2007; The Mathworks Inc, 2018)). Participants performed a visually cued 0.5 Hz (1 second contraction, 1 second rest) repeated force-matching task at either 5% or 50% of their MVC, depending on the session. The MVC was tested immediately prior to executing the submaximal handgrip task in the MRI. The target force was displayed halfway up the vertical axis, and participants were instructed to squeeze the dynamometer to guide a cursor up the screen until it reached the target. During scanning, participants performed 270 handgrip contractions over the nine-minute task.

#### Handgrip force-matching task analysis

The area under the curve (AUC) of the contraction force was calculated over a two second window for every contraction separately. A regression line was then fitted to the AUC measures across all contractions for each participant and session separately. The β of the regression from each session was used to quantify motor performance, whereby a β < 0 indicates a decrement in motor performance over time, consistent with fatigue.

### Response time task

Participants performed a visually-cued response time task before and after MRI scanning, implemented in PsychoPy3 (Peirce et al., 2019). Participants responded via a button box (4-Button Inline, HHSC-1×4-L; Current Designs Inc. Philadelphia, PA USA) with their right or left hand depending on the block. The task was divided into blocks of 64 self-paced visual cues on a computer monitor in a random order, which participants were asked to respond to by pressing the corresponding button on the button box as quickly and accurately as possible. With each hand separately, participants performed two blocks at the Pre-MRI and Post-MRI timepoints (figure 1B).

#### Response time analysis

Incorrect button-press responses, and response times (RTs) < 50 ms or > 700 ms were removed (a total of 656 correct presses were removed across all participants and sessions). This approach to exclude invalid RTs is similar to previous literature (Stagg *et al*., 2011*a*; Steel *et al*., 2016, 2020). The median response time of the remaining button presses was then extrapolated for each block separately. To investigate changes in RT due to repeated handgrip contractions, the percent change in RTs for each hand was calculated using the mean of the medians from the two Pre-MRI blocks as the Pre-MRI value, and similarly, the mean of the medians from the two Post-MRI blocks was used as the Post-MRI value. To investigate any significant change in RT between Pre-MRI and Post-MRI, we performed a RM-ANOVA using the percent change in RT data with one factor of condition (5% MVC, 50% MVC) and another of hand (Left-hand, right-hand).

### Magnetic Resonance Imaging

MRI sessions were acquired using a Siemens 3-tesla Prisma whole-body MRI scanner with a 32-channel head array receive coil (Siemens, Erlangen, Germany). T1-weighted MPRAGE scans (voxel size 1 mm^3^, TR = 1900 ms, TI = 912 ms TE = 3.96 ms, FOV = 232 × 256 × 192 mm^3^, flip angle = 8°, TA = 7:21) were acquired for registration purposes and to guide placement of the MRSI slab, prioritising left M1. Pre- and Post-task resting-state semi-LASER localised density-weighted concentric-ring trajectory MRSI (voxel size 5 mm × 5 mm × 15 mm, semi-LASER VoI 85 mm x 35 mm x 15 mm, TR = 1400 ms, TE = 32 ms, FOV = 240 mm × 240 mm × 15 mm, TA = 4:30) (Steel *et al*., 2018), and fMRI (490 volumes, voxel size 2.4 mm^3^ with zero gap, TR = 735 ms, TE = 39 ms, FOV = 210 × 210 × 154 mm, flip angle = 52°, TA = 6:10) (Miller *et al*., 2016) data were acquired. Participants were asked to fixate on a grey cross presented on a black screen and to blink normally. To quantify any potential change in head position after task performance a single localiser (voxel size 1.4 mm, TR = 3.15 ms, TE = 1.37 ms, FOV = 350 mm × 350 mm × 263 mm, 128 slices, flip angle = 8°, TA = 0:22) was collected prior to Post-task MRSI. Finally, B_0_ field maps (voxel size 2 mm_3_, TR = 482 ms, TE1 = 4.92 ms, TE2 = 7.38 ms, FOV = 216 mm × 216 mm × 146 mm, 49 slices, flip angle = 46°, TA = 1:45) were acquired to correct for field distortion in the resting-state fMRI data.

### MRI analysis

#### fMRI preprocessing

Functional MRI analyses were performed using tools from the FMRIB Software Library (FSL v6.0.3; (Jenkinson *et al*., 2012)). Standard preprocessing steps were performed, including removal of non-brain tissue (BET; (Smith, 2002)), removal of the initial two volumes, motion correction (MCFLIRT; (Jenkinson *et al*., 2002)), high-pass temporal filtering at 0.01Hz, and distortion correction with the implementation of field-maps.

Following single-participant MELODIC pre-processing (v3.15), FMRIB’s Independent Component Analysis (ICA)-based Xnoiseifier (ICA-FIX) was used to automatically denoise the data (Griffanti *et al*., 2014; Salimi-Khorshidi *et al*., 2014). The UK Biobank training-weights file (UKBiobank.RData) was used with a threshold value of 20, and 0.01Hz high-pass filtered motion confound cleanup. Following the automated denoising, all components were manually inspected before the cleaned data were smoothed with a 5 mm full-width half maximum (FWHM) kernel.

Individual resting state fMRI scans were first registered to the respective T1 structural scan using boundary-based registration as implemented in FMRIB’s Linear Image Registration Tool (FLIRT; (Jenkinson & Smith, 2001; Jenkinson *et al*., 2002)), and then to a standard space template (MNI152 2 mm) using non-linear registration (FNIRT; (Andersson *et al*., 2007*a*, 2007*b*)).

#### fMRI analysis

We assessed resting state functional connectivity using two different approaches: (1) a seed-based functional connectivity approach to test the hypothesis that unimanual handgrip contractions lead to changes in connectivity of the ipsilateral right M1 (rM1); and (2) a region of interest (ROI)-based functional connectivity analysis to specifically address our *a priori* hypothesis that unimanual handgrip contractions would increase M1-M1 and SMA-SMA connectivity.

##### (1) Seed-based, whole-brain functional connectivity

The hand area of the rM1 (corresponding to non-contracting left hand) was defined from previous functional MRI data of hand movements (Weinrich *et al*., 2017), and the mean timeseries was extracted from this region for each participant and each session independently. This was then entered as a regressor into a lower-level FEAT analysis (Woolrich *et al*., 2001), and task-related changes in functional connectivity of rM1 were investigated via a higher level mixed-effects analysis using a cluster forming threshold of z = 3.1 and *p* = 0.05 (Woolrich *et al*., 2004).

##### (2) ROI-ROI-based functional connectivity

Next, we wanted to specifically assess changes in M1-M1 and SMA-SMA functional connectivity. The hand area of the M1s was functionally defined as above, and SMA was defined from tractography-based parcellation (Johansen-Berg *et al*., 2004). The mean timeseries from within each region was extracted for each participant and session separately. We calculated the Pearson’s r values between the homologous pairs using custom in-house MATLAB scripts, which were then converted to z-scores using a Fishers *r* to z transformation. The z-scores for the correlation between the homologous pairs were then used in a 2 × 2 × 2 (condition (5%, 50% MVC) × ROI (M1, SMA) × time (pre-task, post-task)) RM-ANOVA.

#### MRSI

MRSI data was reconstructed and pre-processed according to Steel et al. (Steel *et al*., 2018), using in-house scripts. Reconstruction and pre-processing included: i. metabolite cycling reconstruction (Emir *et al*., 2017), ii. coil-combination (Walsh *et al*., 2000), iii. frequency and iv. phase shift correction, v. HLSVD for residual water removal (Cabanes *et al*., 2001), and vi. eddy current correction using the unsuppressed water signal (Klose, 1990). Concentration of neurochemicals was quantified as in Steel et al. (Steel *et al*., 2018) using LCModel (Provencher, 2001, 2016). A chemical shift of 0.5 to 4.2 ppm was evaluated with a basis set containing 20 metabolites and default LCModel macromolecules with disabled soft constraints on metabolites (NRATIO set to 0), and a baseline stiffness setting (DKMTM) of 0.25 (see Figure 4A for example of raw spectra and model fits). Each voxel spectrum was independently fit with spectral quality and assessment of model fitting performed using in-house scrips. For the metabolite maps of glutamate + glutamine (Glx) and GABA, MRSI voxels with CRLB values > 50 or SNR < 40 were excluded from further analysis. All metabolite measurements are expressed as a ratio over total Creatine (tCr).

**Figure 2.**
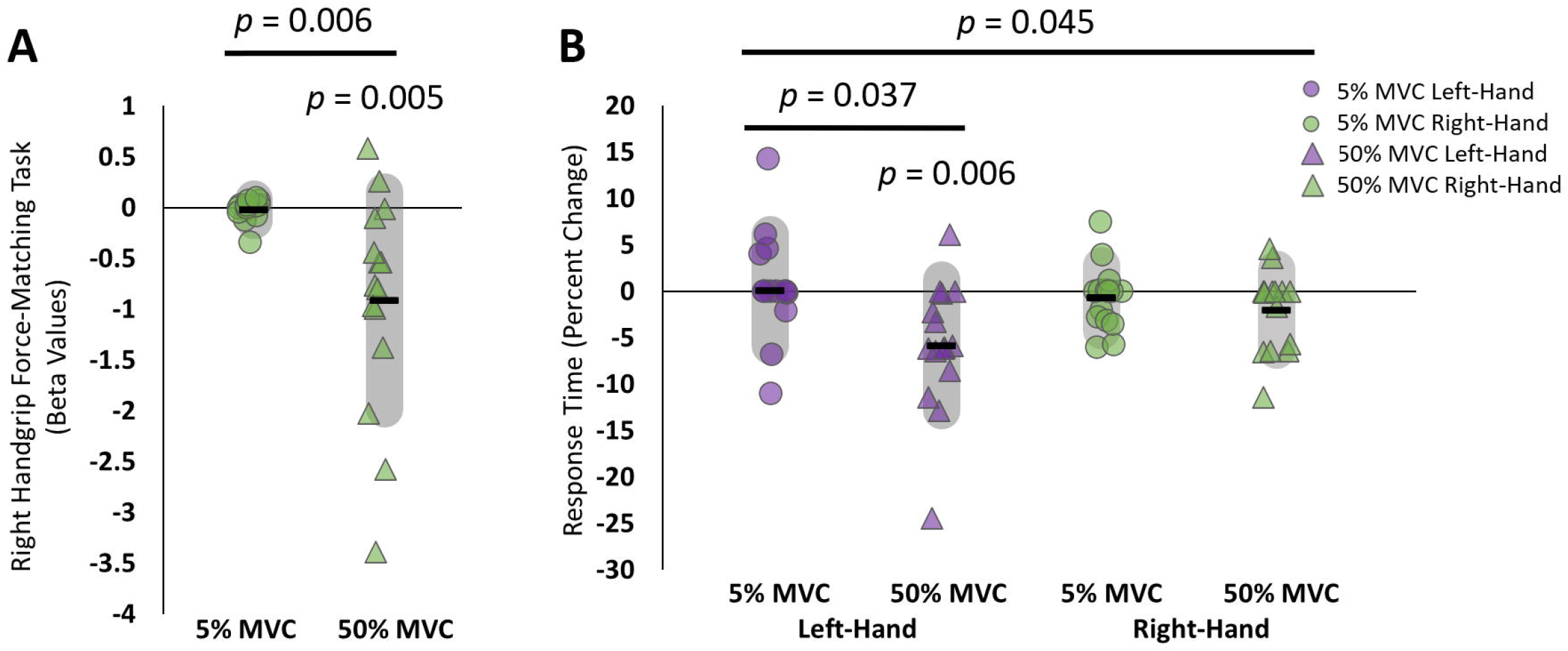
Behavioural changes. **A)** Beta values for the area under the curve of the force profile from the nine-minute right handgrip force-matching task in experiment one. *p* = 0.005: 50% MVC experienced a significant decline in motor performance (one-sample t-test), *p* = 0.006: a significant difference between conditions (paired-samples t-test), and **B)** Percent change in response times from Pre-MRI to Post-MRI for the left (purple data points) and right (green data points) hands after the right-hand force-matching task at 5% MVC (circles) and 50% MVC (triangles). *p* = 0.045: significant condition × hand interaction, *p* = 0.037: significant paired sample t-test, *p* = 0.009: significant one-sample t-test change in response times. Schematic of study design was created with BioRender.com.

**Figure 3.**
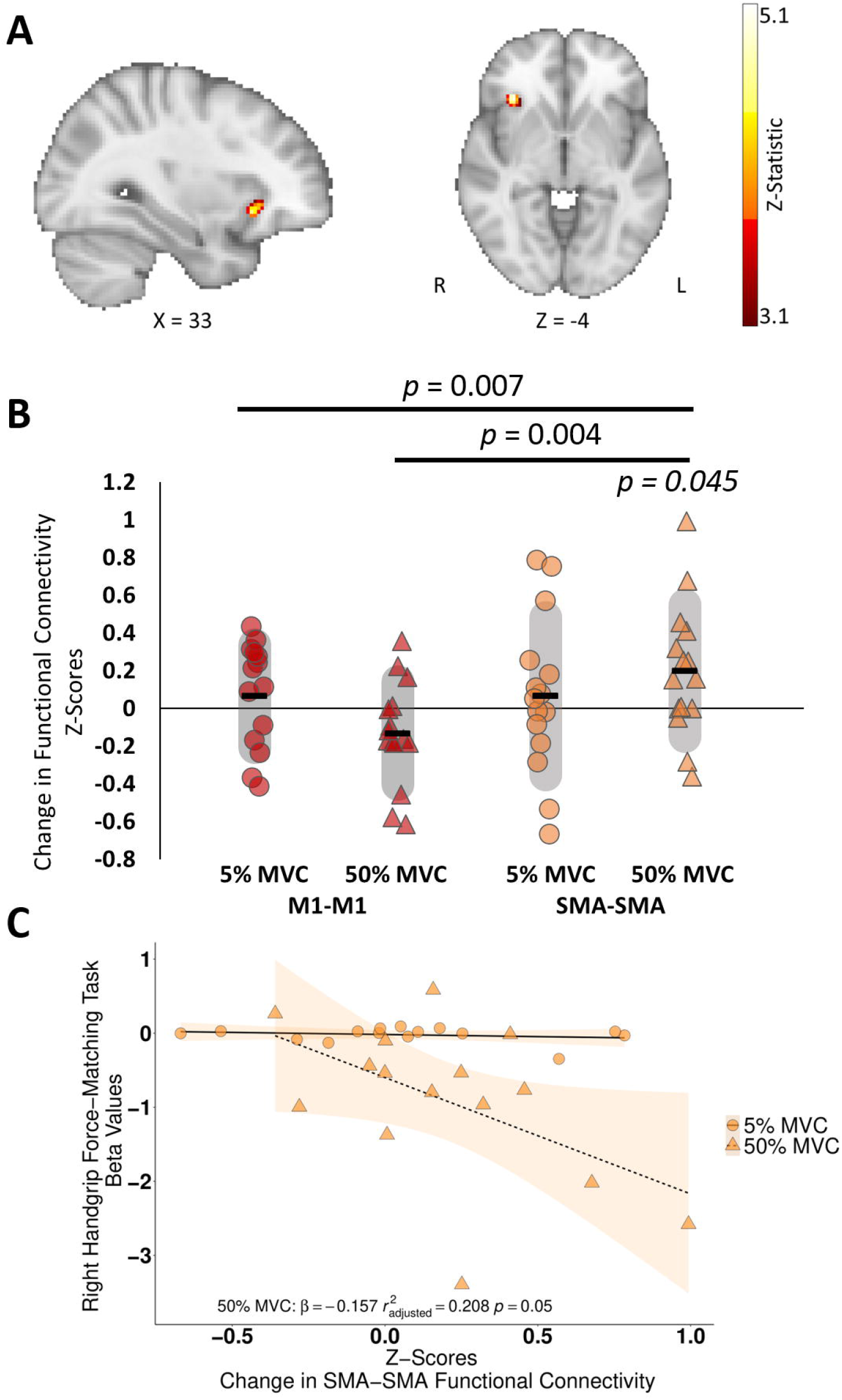
Functional connectivity analyses for **A)** Seed-based functional connectivity analysis between the right ipsilateral primary motor cortex (seed) and the rest of the brain in the 50% MVC condition, data is a Z-statistic threshold map in standard space. Z-threshold = 3.1. Figure is in radiological view (left on right, right on left). **B)** Change in interhemispheric functional connectivity between contralateral and ipsilateral primary motor cortices (M1-M1; red data points), and the contralateral and ipsilateral supplementary motor areas (SMA-SMA; orange data points) for 5% MVC (circles) and 50% MVC (triangles). Back horizonal bar represents the mean change. Grey shaded area represents the standard deviation. *p* = 0.007: significant condition × ROI × time interaction, *p* = 0.004: significant ROI × time interaction for 50% MVC condition, *p* = 0.045: significant paired-sample t-test for SMA-SMA connectivity, and **C)** Scatter plot correlation between X-axis: The change in interhemispheric functional connectivity z-scores between the contralateral left and ipsilateral right supplementary motor areas (SMA-SMA) and Y-axis: beta values representing the decline in motor performance over time (negative slope = performance decline over time) (*r*^2^_adjusted_ = 0.208, *p* = 0.050, β = -0.157). Schematic of study design was created with BioRender.com.

**Figure 4.**
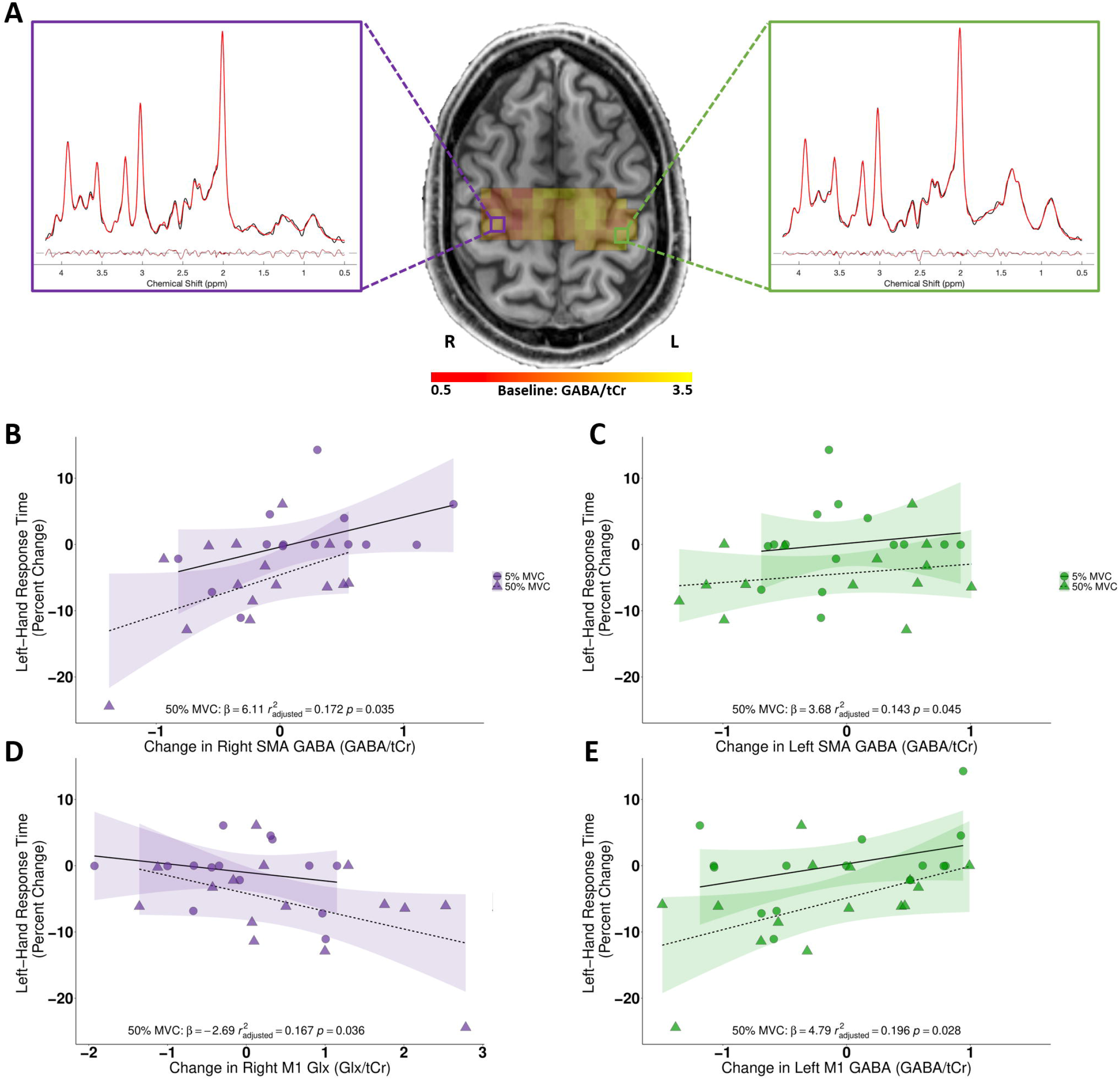
**A)** Magnetic resonance spectroscopic imaging spectrum from a representative participant, with spectra from M1 voxels in each hemisphere. **B)** Scatter plot correlations for the right hemisphere (purple) and left hemisphere (green) between X-axes: SMA (top scatter plots), M1 (bottom scatter plots) for GABA/tCr (rSMA, lSMA, lM1) and Glx/tCr (rM1) ratios with Y-axes: the change in left-hand response times (ms) after performing the right-hand target matching task at either 5% MVC (circles) or 50% MVC (triangles) conditions. Schematic of study design was created with BioRender.com.

Metabolite maps were next upsampled to 1mm^3^ resolution using nearest neighbour interpolation which preserves the original sampling grid. Metabolite maps were then aligned to their native T1 coordinate space. To correct for potential shift in head position, structural scans acquired prior to each MRSI sequence were aligned using McFLIRT (Jenkinson et al., 2002). The metabolite maps were then transformed using the McFLIRT generated registration matrix. Only shared MRSI voxels from within an intersection mask of QA passed and head position aligned MRSI voxels across each timepoint were included in subsequent analyses. For the ROI analysis, the four binary ROI masks [left M1 (lM1), right (rM1), left SMA (lSMA) and right SMA (rSMA)] were non-linearly aligned to native T1 space by inverse warping (3dNwarpApply) the native to standard space alignment, generated with @SSwarper (Cox, 1996; Saad et al., 2009; Cox & Glen, 2013). The native space ROIs were thresholded at 0.5 to mitigate partial volume effects. Within the intersection mask, mean concentration of Glx/tCr and GABA/tCr were then extracted for each of the four ROIs.

### Statistical Analyses

Statistical analyses were run using Jamovi v1.6.9 (The Jamovi Project, 2020). An alpha level for significance testing was set to 0.05, Hedges’ g effect sizes are reported for t-tests and partial eta squared (η_p_^2^) effect sizes are reported for ANOVA results. Greenhouse-Geisser corrections were applied as necessary where violations of sphericity were present. Shapiro-Wilk tests were used to assess data normality. Additionally, the order of sessions was first included in each of the RM-ANOVA models as a covariate, but after determining that the order of testing was not a significant covariate in each of the models (i.e., it did not significantly adjust the dependent variables due to between group differences) it was not included in the final analyses.

## Results

### Performance on the 50% MVC handgrip force-matching task decreased over time

We first wanted to determine if the 50% MVC handgrip force-matching task induced a decrease in handgrip force during task performance. We reasoned that the grip force would be reduced over time during our 50% MVC condition, but not during our 5% MVC control condition, consistent with fatigue. We therefore quantified a line of best fit for each participant for each session. In line with our hypothesis, 50% MVC showed a significant decrement in performance over time compared with 5% MVC (50% MVC: β = -0.912 ± 1.070; 5% MVC: β = -0.022 ± 0.107; slope of 50% MVC compared with zero: *t*(14)= -3.302, *p* = 0.005, g = -0.824; paired t-test *t*(14)= 3.211, *p* = 0.006, g = 0.801; figure 2A).

### Right handgrip contractions decreased RTs in the left, contralateral hand

We next investigated whether performance of a 50% MVC force-matching task with the right hand would induce a behavioural improvement of the left hand. After running a condition × hand RM-ANOVA with the RT percent change data, we observed a significant condition × hand interaction (*F*(1,14)= 4.84, *p* = 0.045, η_p_^2^ = 0.257), but no significant main effects. To explore the interaction, we ran paired sample t-tests between conditions for each hand separately. This revealed a significant decrease in RTs in the left hand for the 50% MVC condition compared with the 5% MVC condition (5%: Δ = 0.08 ± 5.99%; 50%: Δ = -5.85 ± 7.05%; *t*(14)= 2.30, *p* = 0.037, g = 0.574). Further, the left hand RT improvement was significantly different from zero (*t*(14)= -3.21, *p* = 0.006, g = -0.801). There were no significant differences for the right hand between the 50% MVC (Δ = -2.02 ± 4.35%) and 5% MVC (Δ = -0.73 ± 3.46%) conditions (*t*(14)= 0.788, *p* = 0.441, g = 0.196; figure 2B).

### Repeated 50% MVC unilateral handgrip contractions increase functional connectivity between the ipsilateral rM1 and right orbitofrontal cortex

To investigate the neural changes associated with our observed behavioural improvements in the left, contralateral hand, we performed a voxel-wise seed-based analysis from the hand area of ipsilateral, rM1. There was a significant increase in functional connectivity with the ipsilateral, right orbitofrontal cortex (rOFC) in the 50% MVC condition, compared with the 5% MVC condition (table 1; figure 3A).

**Table 1.** Post > Pre-force-matching task rs-fMRI contrasts from iM1 seed-based functional connectivity analysis.

| 50% MVC |  |  |  |  |  |  |
| --- | --- | --- | --- | --- | --- | --- |
| MNI Coordinates (mm) |  |  |  |  |  |  |
| Voxels | *p*-value | z-max | X | Y | Z | Label |
| 104 | 0.0348 | 5.12 | 34 | 26 | -6 | Right Orbitofrontal Cortex |
| 5% MVC |  |  |  |  |  |  |
| MNI Coordinates (mm) |  |  |  |  |  |  |
| Voxels | *p*-value | z-max | X | Y | Z | Label |
| 182 | 0.00187 | 4.44 | 28 | -50 | -26 | Right Cerebellum VI |
| 104 | 0.0363 | 4.78 | -42 | -54 | -4 | Left Inferior Temporal Gyrus |

### SMA-SMA connectivity increased after 50% MVC contractions

To address our *a priori* hypothesis that the 50% MVC condition would alter inter-hemispheric functional connectivity between left and right M1, and between left and right SMA, we performed an ROI-ROI connectivity analysis. A RM-ANOVA with one factor of condition (5% MVC, 50% MVC), one factor of ROI (M1, SMA), and one factor of time (Pre-Task, Post-Task) revealed a significant three-way (condition × ROI × time) interaction [(*F*(1,14) = 10.154, *p* = 0.007, η_p_^2^ = 0.420); a significant main effect of ROI (*F*(1,14) = 118.113, *p* < 0.001, η_p_^2^ = 0.894), but no significant main effect of condition (*F*(1,14) = 0.508, *p* = 0.488, η_p_^2^ = 0.8035), or time (*F*(1,14) = 0.797, *p* = 0.387, η_p_^2^ = 0.054)].

To understand this three-way interaction, we then ran separate RM-ANOVA tests for the 50% and 5% MVC conditions, which revealed a significant ROI × time interaction for the 50% MVC condition (*F*(1,14) = 11.970, *p* = 0.004, η_p_^2^ = 0.461) but not for the 5% MVC condition (*F*(1,14) < 0.001, *p* = 0.997, η_p_^2^ < 0.001). Follow-up tests revealed a significant increase in SMA-SMA connectivity in the 50% MVC condition (*t*(14) = -2.203, *p* = 0.045, g = -0.550), figure 3B), but no significant changes in the 5% MVC, nor for the M1 in either condition.

Given the 50% MVC force-matching task led to both a decrease in force output and an increase in SMA-SMA connectivity, we next wanted to investigate whether these two effects were related. We demonstrated a negative correlation between the degree of force decline in the 50% MVC condition and change in SMA-SMA connectivity, such that a greater decline in handgrip performance, indexed by greater decrease in the force output, correlated with an increase in SMA-SMA connectivity (*r*^*2*^_adjusted_ = 0.208, *p* = 0.050, β = -0.157; figure 3C). This relationship was not observed for M1-M1 connectivity (50% MVC: M1-M1 connectivity and force decline *r*^2^_adjusted_ = -0.077, *p* = 0.945, β = -0.076) or in SMA-SMA connectivity in the 5% MVC condition (5% MVC: SMA-SMA connectivity and force decline: *r*^2^_adjusted_ = -0.024, *p* = 0.427, β = - 0.863).

### Decrease in SMA and M1 GABA correlated with decrease in left-hand RTs after right-hand 50% MVC handgrip contractions

Finally, given our *a priori* hypothesis that the behavioural effects of unilateral 50% MVC handgrip contractions decrease inhibition, we investigated changes in GABA within each ROI. We ran RM-ANOVA tests for M1 and SMA separately, with one factor of condition (50% MVC, 5% MVC), one factor of hemisphere (left, right) and one factor of time (pre, post). There were no significant main effects or interactions. However, in line with our *a priori* hypothesis that GABA would decrease and Glx would increase in response to handgrip contractions, there were significant positive correlations between an individual participant’s decrease in RTs for the left-hand and their decrease in lM1 GABA (one-tailed: *r*^*2*^_adjusted_ = 0.196, *p* = 0.028, β = 4.79), lSMA GABA (one-tailed: *r*^*2*^_adjusted_ = 0.143, *p* = 0.045, β = 3.68), and rSMA GABA (one-tailed: *r*^*2*^_adjusted_ = 0.172, *p* = 0.035, β =6.11) such that the greater the decrease in GABA in these regions, the greater behavioural improvement. A significant negative correlation was also observed for the increase in rM1 Glx with the decrease in RTs for the left-hand (one-tailed: *r*^*2*^_adjusted_ = 0.167, *p* = 0.036, β = -2.69; figure 4).

## Discussion

The objective of this study was to determine the neural correlates of unimanual 50% MVC handgrip contractions and the impact on contralateral limb motor performance. Specifically, we hypothesised that a decrease in handgrip force would be associated with increased sensorimotor functional connectivity (Jiang *et al*., 2012) and disinhibition of ipsilateral motor areas (Takahashi *et al*., 2009), neural effects that lead to enhanced performance of the contralateral, non-contracting hand.

We showed that right handgrip contractions at 50% MVC, but not 5% MVC, resulted in a decline in handgrip force, something that would be consistent with fatigue, though other interpretations are possible. We also observed, in line with our hypothesis, that repeated 50% MVC handgrip contractions performed with the right-hand led to significant improvements in left hand RTs, something that was not seen either in the right hand, nor with either hand in the low force 5% MVC condition. These improvements were also accompanied by neural changes. We observed a significant interhemispheric interaction between a decline in M1-M1 connectivity and an increase in SMA-SMA connectivity. The increase in SMA-SMA connectivity correlated with the degree of force decline on a participant-by-participant basis. Additionally, increased rM1-rOFC connectivity was observed on a voxel-wise analysis. Finally, we used MRSI to quantify potential changes in GABA and Glx. We observed no group mean changes, but on a participant-by-participant basis, GABA decreases in the lM1 and bilateral SMAs modestly correlated with behavioural improvements.

### Increased functional connectivity between ipsilateral rM1 and rOFC after repeated right 50% MVC handgrip contractions

We observed increased functional connectivity between our rM1 seed and the rOFC after right 50% MVC handgrip contractions. Increased OFC activity has been associated with faster response times (Bode *et al*., 2018) and hand motor learning (Alves Heinze *et al*., 2019). The increased rM1-rOFC functional connectivity observed here may therefore reflect the OFC’s role as a top-down motor control region (Ono *et al*., 2014) which is involved in the regulation of motor responses and error monitoring (Alves Heinze *et al*., 2019). In line with this hypothesis, Jackson et al. (2003) showed increased activation in the right OFC in participants who significantly improved their performance in a right-hand motor task, either having physically executed the task or having performed motor imagery. Taken together, our results suggest that increased functional connectivity between the rM1 and the rOFC may be functionally relevant for improved RTs in the contralateral, non-contracting left-hand.

### Repetitive 50% MVC contractions increase SMA-SMA connectivity

Our ROI-based functional connectivity analysis examined how the temporal correlations between left and right M1 and SMA homologs changed after the right handgrip force-matching task. After 50% MVC contractions, SMA-SMA functional connectivity increased, and the degree of change was negatively correlated with the degree of decreased handgrip force, as indexed by a decrease in the area-under-the-curve for each contraction over time, such that participants who exhibited greater declines in this metric showed greater increase in SMA-SMA functional connectivity.

The modulation of the ipsilateral SMA after unilateral fatigue may be explained in part due to its dense interhemispheric homologous connections (Ruddy *et al*., 2017) and is thought to play an important role in interhemispheric communication in the movement preparation phase (Welniarz *et al*., 2019). The inter-connectedness of the SMA between hemispheres offers insight as to why unilateral fatiguing exercise modulates the ipsilateral motor network. Further, lesion studies have demonstrated that impairments to the SMA cause increased response time (i.e., slower movement) (Viallet *et al*., 1995).

The SMA has been described as a ‘phylogenetically older’ M1 (Goldberg, 1985) which has direct excitatory connections not only onto the M1 circuitry (Shirota *et al*., 2012; Arai *et al*., 2012), but onto alpha-motoneurons that innervate the hand and fingers (Maier *et al*., 2002; Dum & Strick, 2005). The SMA plays a major role in the planning and self-initiation of voluntary movement, and serves to integrate multimodal information to ensure motor and visual systems are in agreement (Hikosaka *et al*., 1999; Nakahara *et al*., 2001). The increased connectivity between left and right SMA in the present study may reflect a mechanism by which repetitive unilateral 50% MVC handgrip contractions induced neural changes that improve motor performance for the inactive, contralateral hand. Therefore, it is plausible that the 50% MVC condition-specific increase in SMA-SMA functional connectivity improved contralateral left-hand RTs either by influencing rM1 through rSMA-rM1 excitatory connections or through direct corticospinal connections to upper limb alpha-motoneurons (Dum & Strick, 2005). Transcallosal homologous connectivity between motor regions primarily reflects inhibitory processes (Wahl *et al*., 2007). Therefore, an increase in SMA-SMA functional connectivity may reflect an increase in interhemispheric inhibition between these premotor regions. Given this, we can hypothesize that an increase in interhemispheric SMA-SMA functional connectivity, reflecting interhemispheric inhibition, may aid in silencing unwanted cortical activity that facilitates a more focal excitatory process in the opposite hemisphere (Carson, 2020) giving rise to enhanced motor performance (i.e., faster RTs) in the opposite hand.

### A decrease in lM1 and bilateral SMA GABA after a decrease in handgrip force correlates with improvements in left-hand RTs

To address our hypothesis that the 50% MVC contractions would lead to decreased inhibition in motor regions, we used a novel MRSI sequence to quantify neurochemical concentrations across the sensorimotor network. At a group level, there were no significant GABA changes in M1 and SMA in either hemisphere after 50% MVC contractions compared with 5% MVC control condition. However, in line with our *a priori* hypothesis, we did observe significant one-tailed, uncorrected correlations on a participant-by-participant basis between the decrease in lM1, lSMA, and rSMA GABA after right 50% MVC contractions and the improved behaviour in the non-contracting hand. While the mechanism by which lM1 and bilateral SMA GABA decreases lead to behavioural improvement in the left contralateral hand after right handgrip contractions is unclear, a combination of decussated and non-decussated pathways may play an important role (Maruyama *et al*., 2006; Aboodarda *et al*., 2016).

## Conclusions

This study identified that repetitive 50% MVC right handgrip contractions led to behavioural improvements in the opposite hand which were accompanied by increased SMA-SMA functional connectivity. In addition, we demonstrated an increase in functional connectivity between rM1 and the rOFC. We also demonstrated positive relationships between change in inhibition (GABA) in lM1 and bilateral SMA and a negative relationship between the change in rM1 Glx with the behavioural improvement in the non-contracting left-hand. These results suggest that repetitive 50% MVC unimanual contractions can enhance subsequent motor performance of the opposite hand. This approach may serve as a promising adjunct therapy to prime the ipsilateral sensorimotor system for rehabilitation in a range of neurological or orthopedic conditions.

## Author contributions

J.W.A., J.M.L, J.P.F., & C.J.S. conceived and designed the experiment. C.Z. & J.M.L developed the software to run the behavioural experiment. J.W.A., & J.M.L. collected the data and J.W.A., E.M., J.M.L., J.P.F., & C.J.S. analyzed the data. All authors were involved in data interpretation. J.W.A. drafted the paper, and all authors revised it and approve the final version.

## Conflict of interest

None to report

## Funding sources

C.J.S. holds a Sir Henry Dale Fellowship, funded by the Wellcome Trust and the Royal Society (102584/Z/13/Z). This work was also supported by the National Institute for Health Research (NIHR) Oxford Health Biomedical Research Centre, and the John Fell Fund. The Wellcome Centre for Integrative Neuroimaging is supported by core funding from the Wellcome Trust (203139/Z/16/Z). J.P.F. holds a Natural Sciences and Engineering Research Council of Canada (NSERC) Discovery Grant: RGPIN 2016-0529 at the University of Saskatchewan. J.M.L was supported by a National Institutes of Health Oxford-Cambridge Scholar Fellowship, and the International Biomedical Research Alliance. J.W.A. received an NSERC Alexander Graham Bell Canada Graduate Scholarship – Doctoral (CGS D) Award and a MITACS Globalink Research Award to complete this work.

## Acknowledgements

This research was funded in whole, or in part, by the Wellcome Trust [Grant number 102584/Z/13/B and is 203139/Z/16/Z]. For the purpose of open access, the author has applied a CC BY public copyright licence to any Author Accepted Manuscript version arising from this submission. The authors would like to acknowledge Dr. Jason DeFreitas for his valuable insight on the manuscript.

## Notes

### Competing Interest Statement

The authors have declared no competing interest.

### Summary of Updates

The original version of this manuscript contained two components essentially serving as two separate experiments. This version of the manuscript has been rewritten and the second experiment has been removed to improve clarity.

## References

Aboodarda SJ, Šambaher N & Behm DG (2016). Unilateral elbow flexion fatigue modulates corticospinal responsiveness in non-fatigued contralateral biceps brachii. Scand J Med Sci Sport 26, 1301–1312.

Aboodarda SJ, Zhang CXY, Sharara R, Cline M & Millet GY (2019). Exercise-induced fatigue in one leg does not impair the neuromuscular performance in the contralateral leg but improves the excitability of the ipsilateral corticospinal pathway. Brain Sci 9, 250.

Allman C, Amadi U, Winkler AM, Wilkins L, Filippini N, Kischka U, Stagg CJ & Johansen-Berg H (2016). Ipsilesional anodal tDCS enhances the functional benefits of rehabilitation in patients after stroke. Sci Transl Med 8, 330re1–330re1.

Alves Heinze R, Vanzella P, Zimeo Morais GA & Sato JR (2019). Hand motor learning in a musical context and prefrontal cortex hemodynamic response: a functional near-infrared spectroscopy (fNIRS) study. Cogn Process; DOI: 10.1007/s10339-019-00925-y.

Andersson JLR, Jenkinson M & Smith S (2007a). Non-linear registration aka spatial normalisation FMRIB Technial Report TR07JA2.

Andersson JLR, Jenkinson M & Smith S (2007b). Non-linear optimisation FMRIB Technial Report TR07JA1.

Arai N, Lu MK, Ugawa Y & Ziemann U (2012). Effective connectivity between human supplementary motor area and primary motor cortex: A paired-coil TMS study. Exp Brain Res 220, 79–87.

Bachtiar V, Johnstone A, Berrington A, Lemke C, Johansen-Berg H, Emir U & Stagg CJ (2018). Modulating regional motor cortical excitability with noninvasive brain stimulation results in neurochemical changes in bilateral motor cortices. J Neurosci 38, 7327–7336.

Bachtiar V, Near J, Johansen-Berg H & Stagg CJ (2015). Modulation of GABA and resting state functional connectivity by transcranial direct current stimulation. Elife 4, e08789.

Bode S, Bennett D, Sewell DK, Paton B, Egan GF, Smith PL & Murawski C (2018). Dissociating neural variability related to stimulus quality and response times in perceptual decisionmaking. Neuropsychologia 111, 190–200.

Branscheidt M, Kassavetis P, Anaya M, Rogers D, Huang HD, Lindquist MA & Celnik P (2019). Fatigue induces long-lasting detrimental changes in motor-skill learning. Elife; DOI: 10.7554/eLife.40578.

Cabanes E, Confort-Gouny S, Le Fur Y, Simond G & Cozzone PJ (2001). Optimization of residual water signal removal by HLSVD on simulated short echo time proton MR spectra of the human brain. J Magn Reson 150, 116–125.

Carr JC, Bemben M, Stock MS & DeFreitas JM (2021). Ipsilateral and contralateral responses following unimanual fatigue with and without illusionary mirror visual feedback. J Neurophysiol jn.00077.2021.

Carson RG (2020). Inter-hemispheric inhibition sculpts the output of neural circuits by co-opting the two cerebral hemispheres. J Physiol J P279793.

Cirillo J, Rogasch NC & Semmler JG (2010). Hemispheric differences in use-dependent corticomotor plasticity in young and old adults. Exp brain Res 205, 57–68.

Cirillo J, Todd G & Semmler JG (2011). Corticomotor excitability and plasticity following complex visuomotor training in young and old adults. Eur J Neurosci 34, 1847–1856.

Cox RW (1996). AFNI: software for analysis and visualization of functional magnetic resonance neuroimages. Comput Biomed Res 29, 162–173.

Cox RW & Glen DR (2013). Nonlinear warping in AFNI. Present 19th Annu Meet Organ Hum Brain Mapp. Available at: https://afni.nimh.nih.gov/pub/dist/HBM2013/Cox_Poster_HBM2013.pdf.

Dum RP & Strick PL (2005). Frontal lobe inputs to the digit representations of the motor areas on the lateral surface of the hemisphere. J Neurosci 25, 1375–1386.

Emir UE, Burns B, Chiew M, Jezzard P & Thomas MA (2017). Non-water-suppressed short-echo-time magnetic resonance spectroscopic imaging using a concentric ring k-space trajectory. Nmr Biomed; DOI: 10.1002/NBM.3714.

Floyer-Lea A, Wylezinska M, Kincses T & Matthews PM (2006). Rapid modulation of GABA concentration in human sensorimotor cortex during motor learning. J Neurophysiol 95, 1639–1644.

Goldberg G (1985). Supplementary motor area structure and function: Review and hypotheses. Behav Brain Sci 8, 567–588.

Griffanti L, Salimi-Khorshidi G, Beckmann CF, Auerbach EJ, Douaud G, Sexton CE, Zsoldos E, Ebmeier KP, Filippini N, Mackay CE, Moeller S, Xu J, Yacoub E, Baselli G, Ugurbil K, Miller KL & Smith SM (2014). ICA-based artefact removal and accelerated fMRI acquisition for improved resting state network imaging. Neuroimage; DOI: 10.1016/j.neuroimage.2014.03.034.

Hikosaka O, Sakai K, Lu X, Nakahara H, Rand MK, Nakamura K, Miyachi S & Doya K (1999). Parallel neural networks for learning sequential procedures. Trends Neurosci 22, 464–471. Available at: http://www.cell.com/article/S0166223699014393/fulltext [Accessed April 15, 2021].

Jackson PL, Lafleur MF, Malouin F, Richards CL & Doyon J (2003). Functional cerebral reorganization following motor sequence learning through mental practice with motor imagery. Neuroimage; DOI: 10.1016/S1053-8119(03)00369-0.

Jenkinson M, Bannister P, Brady M & Smith S (2002). Improved optimization for the robust and accurate linear registration and motion correction of brain images. Neuroimage 17, 825–841.

Jenkinson M, Beckmann CF, Behrens TEJ, Woolrich MW & Smith SM (2012). FSL. Neuroimage 62, 782–790.

Jenkinson M & Smith S (2001). A global optimisation method for robust affine registration of brain images. Med Image Anal 5, 143–156.

Jiang Z, Wang X-F, Kisiel-Sajewicz K, Yan JH & Yue GH (2012). Strengthened functional connectivity in the brain during muscle fatigue. Neuroimage 60, 728–737.

Johansen-Berg H, Behrens TEJ, Robson MD, Drobnjak I, Rushworth MFS, Brady JM, Smith SM, Higham DJ & Matthews PM (2004). Changes in connectivity profiles define functionally distinct regions in human medial frontal cortex. Proc Natl Acad Sci U S A 101, 13335–13340.

Kavanagh JJ, Feldman MR & Simmonds MJ (2016). Maximal intermittent contractions of the first dorsal interosseous inhibits voluntary activation of the contralateral homologous muscle. J Neurophysiol 116, 2272–2280.

Kim D-Y, Lim J-Y, Kang EK, You DS, Oh M-K, Oh B-M & Paik N-J (2010). Effect of transcranial direct current stimulation on motor recovery in patients with subacute stroke. Am J Phys Med Rehabil 89, 879–886.

Kleiner M, Brainard DH, Pelli DG, Broussard C, Wolf T & Niehorster D (2007). What’s new in Psychtoolbox-3? Perception.

Klose U (1990). In vivo proton spectroscopy in presence of eddy currents. Magn Reson Med 14, 26–30.

Kolasinski J, Hinson EL, Divanbeighi Zand AP, Rizov A, Emir UE & Stagg CJ (2019). The dynamics of cortical GABA in human motor learning. J Physiol; DOI: 10.1113/JP276626.

Lissek S, Vallana GS, Güntürkün O, Dinse H & Tegenthoff M (2013). Brain activation in motor sequence learning Is related to the level of native cortical excitability. PLoS One 8, e61863.

Maier MA, Armand J, Kirkwood PA, Yang H-W, Davis JN & Lemon R.N. (2002). Differences in the corticospinal projection from primary motor cortex and supplementary motor area to macaque upper limb motoneurons: An anatomical and electrophysiological study. Cereb Cortex 12, 281–296.

Maruyama A, Matsunaga K, Tanaka N & Rothwell JC (2006). Muscle fatigue decreases short-interval intracortical inhibition after exhaustive intermittent tasks. Clin Neurophysiol 117, 864–870.

Miller KL et al. (2016). Multimodal population brain imaging in the UK Biobank prospective epidemiological study. Nat Neurosci 2016 1911 19, 1523–1536.

Nakahara H, Doya K & Hikosaka O (2001). Parallel cortico-basal ganglia mechanisms for acquisition and execution of visuomotor sequences - A computational approach. J Cogn Neurosci 13, 626–647.

Oldfield RC (1971). The assessment and analysis of handedness: The Edinburgh inventory. Neuropsychologia 9, 97–113.

Ono Y, Nomoto Y, Tanaka S, Sato K, Shimada S, Tachibana A, Bronner S & Noah JA (2014). Frontotemporal oxyhemoglobin dynamics predict performance accuracy of dance simulation gameplay: Temporal characteristics of top-down and bottom-up cortical activities. Neuroimage 85, 461–470.

Peirce J, Gray JR, Simpson S, MacAskill M, Höchenberger R, Sogo H, Kastman E & Lindeløv JK (2019). PsychoPy2: Experiments in behavior made easy. Behav Res Methods 51, 195–203.

Provencher SW (2001). Automatic quantitation of localized in vivo 1H spectra with LCModel. NMR Biomed 14, 260–264.

Provencher SW (2016). LCModel and LCMgui User’s Manual. Available at: http://s-provencher.com/lcm-manual.shtml.

Reis J, Schambra HM, Cohen LG, Buch ER, Fritsch B, Zarahn E, Celnik PA & Krakauer JW (2009). Noninvasive cortical stimulation enhances motor skill acquisition over multiple days through an effect on consolidation. Proc Natl Acad Sci U S A 106, 1590–1595.

Ruddy KL, Leemans A & Carson RG (2017). Transcallosal connectivity of the human cortical motor network. Brain Struct Funct 222, 1243–1252.

Saad ZS, Glen DR, Chen G, Beauchamp MS, Desai R & Cox RW (2009). A New Method for Improving Functional-to-Structural MRI Alignment using Local Pearson Correlation. Neuroimage 44, 839.

Salimi-Khorshidi G, Douaud G, Beckmann CF, Glasser MF, Griffanti L & Smith SM (2014). Automatic denoising of functional MRI data: Combining independent component analysis and hierarchical fusion of classifiers. Neuroimage 90, 449–468.

Sanes JN & Donoghue JP (2000). Plasticity and primary motor cortex. Annu Rev Neurosci 23, 393–415.

Shirota Y, Hamada M, Terao Y, Ohminami S, Tsutsumi R, Ugawa Y & Hanajima R (2012). Increased primary motor cortical excitability by a single-pulse transcranial magnetic stimulation over the supplementary motor area. Exp Brain Res 219, 339–349.

Smith SM (2002). Fast robust automated brain extraction. Hum Brain Mapp 17, 143–155.

Stagg CJ, Bachtiar V, Amadi U, Gudberg CA, Ilie AS, Sampaio-Baptista C, O’Shea J, Woolrich M, Smith SM, Filippini N, Near J & Johansen-Berg H (2014). Local GABA concentration is related to network-level resting functional connectivity. Elife 3, 1–9.

Stagg CJ, Bachtiar V & Johansen-Berg H (2011a). The role of GABA in human motor learning. Curr Biol 21, 480–484.

Stagg CJ, Jayaram G, Pastor D, Kincses ZT, Matthews PM & Johansen-Berg H (2011b). Polarity and timing-dependent effects of transcranial direct current stimulation in explicit motor learning. Neuropsychologia 49, 800–804.

Steel A, Baker CI & Stagg CJ (2020). Intention to learn modulates the impact of reward and punishment on sequence learning. Sci Reports 2020 101 10, 1–13.

Steel A, Chiew M, Jezzard P, Voets NL, Plaha P, Thomas MA, Stagg CJ & Emir UE (2018). Metabolite-cycled density-weighted concentric rings k-space trajectory (DW-CRT) enables high-resolution 1 H magnetic resonance spectroscopic imaging at 3-Tesla. Sci Rep 8, 7792.

Steel A, Silson EH, Stagg CJ & Baker CI (2016). The impact of reward and punishment on skill learning depends on task demands. Sci Reports 2016 61 6, 1–9.

Takahashi K, Maruyama A, Maeda M, Etoh S, Hirakoba K, Kawahira K & Rothwell JC (2009). Unilateral grip fatigue reduces short interval intracortical inhibition in ipsilateral primary motor cortex. Clin Neurophysiol 120, 198–203.

The Jamovi Project (2020). Jamovi (Version 1.2). Available at: https://www.jamovi.org.

The Mathworks Inc (2018). MATLAB 2018a. http://www.MathworksCom/Products/Matlab.

Veale JF (2014). Edinburgh Handedness Inventory - Short Form: A revised version based on confirmatory factor analysis. Laterality 19, 164–177.

Viallet F, Vuillon-Cacciuttolo G, Legallet E, Bonnefoi-Kyriacou B & Trouche E (1995). Bilateral and side-related reaction time impairments in patients with unilateral cerebral lesions of a medial frontal region involving the supplementary motor area. Neuropsychologia 33, 215–223.

Wahl M, Lauterbach-Soon B, Hattingen E, Jung P, Singer O, Volz S, Klein JC, Steinmetz H & Ziemann U (2007). Human motor corpus callosum: Topography, somatotopy, and link between microstructure and function. J Neurosci 27, 12132–12138.

Walsh DO, Gmitro AF & Marcellin MW (2000). Adaptive reconstruction of phased array MR imagery. Magn Reson Med; DOI: 10.1002/(SICI)1522-2594(200005)43:5.

Weinrich CA, Brittain JS, Nowak M, Salimi-Khorshidi R, Brown P & Stagg CJ (2017). Modulation of long-range connectivity patterns via frequency-specific stimulation of human cortex. Curr Biol 27, 3061–3068.e3.

Welniarz Q et al. (2019). The supplementary motor area modulates interhemispheric interactions during movement preparation. Hum Brain Mapp 40, 2125.

Woolrich MW, Behrens TEJ, Beckmann CF, Jenkinson M & Smith SM (2004). Multilevel linear modelling for FMRI group analysis using Bayesian inference. Neuroimage 21, 1732–1747.

Woolrich MW, Ripley BD, Brady M & Smith SM (2001). Temporal autocorrelation in univariate linear modeling of FMRI data. Neuroimage 14, 1370–1386.

